## Supporting Material for "A methodology to reduce the localization error in multi-loci microscopy provides new insights into enhancer biology"

### Supporting Material for A methodology to reduce the localization error of multi-loci microscopy data provides new insights into enhancer biology

**Christopher H. Bohrer and Daniel R. Larson***

*Laboratory of Receptor Biology and Gene Expression, Center for Cancer Research, National Cancer Institute, National Institutes of Health, Bethesda, Maryland 20892, USA*

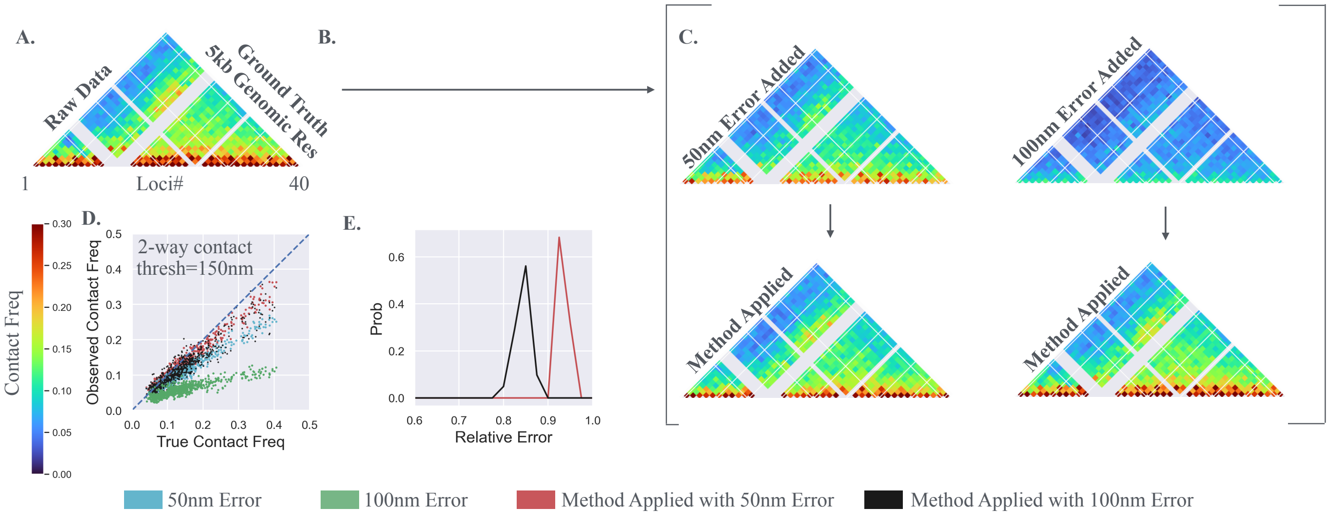


Figure S1: *Testing the Methodology with Simulation Raw Data.* These results are similar to Main Fig. 2. A) The contact frequencies using a distance threshold of 150nm for the chromatin tracing data of Huang et al. (4) (no boundary condition). B) We do not apply our methodology and just computationally add the localization error. That is, we use the raw empirical locations as a ground truth. C) We add localization error directly to the raw data and generate the contact frequencies for the labeled localization error. In the lower row the contact frequencies that result from the application of the methodology are shown. D) The observed/quantified contact frequencies vs. the true underlying contact frequencies, again quantified using the raw data. E) The relative amount of localization error quantified for individual cells, see main text.

### Methodology to Maximize Likelihood (Equation #3 Main Text)

To determine the best estimation of $A^{\ell}$ (see main text) we utilized a simple stochastic descent algorithm (see below for pseudocode). While we initially tried to apply a Markov Chain Monte Carlo approach, we found that utilizing a more simple algorithm converged much quicker and resulted in a better performance given a 'reasonable' computation time (data not shown). We believe this to be the result of the high dimensionality of the problem, and, similarly, the fact that the more simple stochastic descent algorithm was more focused around the initial observed locations. That is, given the error in quantifying the likelihood due to some of our approximations (discussed more later), it was better to find the local minimum 'closest' to the starting positions of the localizations. We note that our investigation of different algorithms was not exhaustive and the application of other methods may prove beneficial in the future.

### Approximating the probability distribution for goal variances

We performed the following to approximate the parameters that describe the probability of observing a particular goal variance; here we show it for a pair of loci $(\alpha, \beta)$ given the empirical data $(O^{\alpha}, O^{\beta})$ and the estimate of the

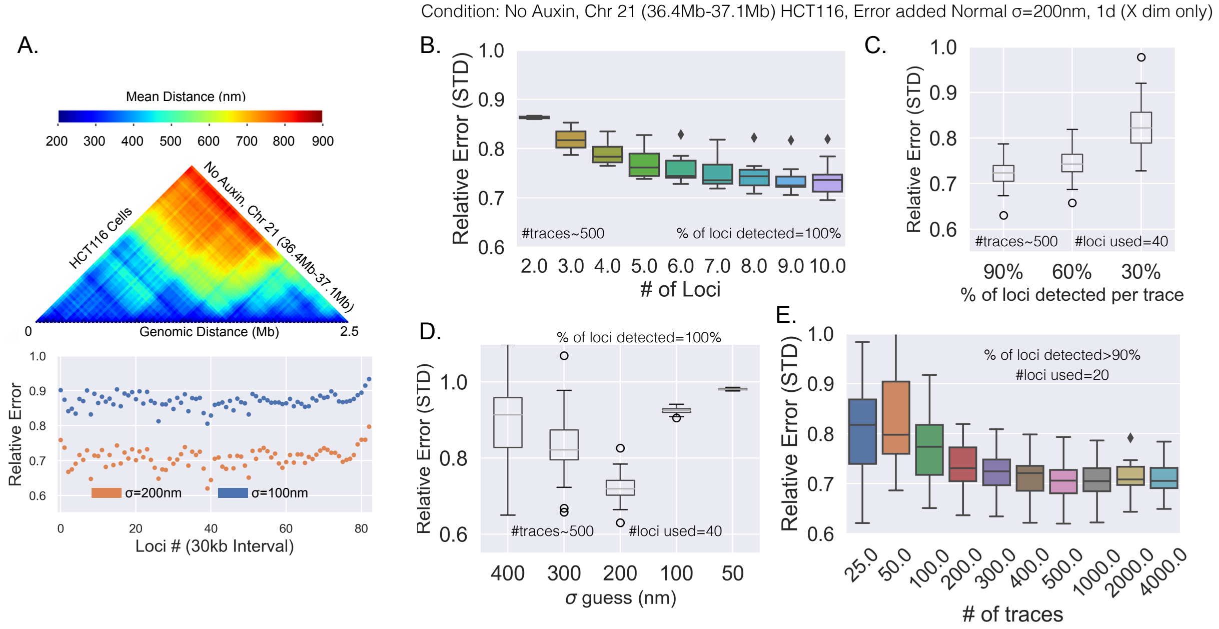


Figure S2: *The effectiveness of the methodology given different parameters:* A) Top: the mean distances for the chromatin tracing data of Bintu et al. (1) for the condition labeled; all analysis within this figure uses this dataset. Bottom: The relative error for each loci after the methodology for the amount of localization error introduced. B) The relative error vs the number of loci included in the application of the methodology. C) The relative error as a function of the detection efficiency. D) The relative error as a function of the localization error guess; the true localization error equals 200nm. E) The relative error vs the number of traces included.

localization error ($Var(\epsilon^\alpha)$).

The goal is to determine the probability distribution of $Var(T^\alpha - O^\beta)$. Because this is a variance quantified with many datapoints, the probability distribution will approach a gaussian distribution due to the central limit theorem. With this, we then need to quantify the mean and variance ($\sigma^2_{\alpha,\beta}$) of this gaussian distribution. The mean is simply: $Var(O^\alpha - O^\beta) - Var(\epsilon^\alpha)$, as described within the main text. The quantification of $\sigma^2_{\alpha,\beta}$ is an approximation, and the specific approach we took is shown in algorithm 2; given all systems investigated within this work, the approach described within algorithm 2 worked relatively well, resulting in approximately 10 to 20 percent error (data not shown). We note that more accurate calculations of $\sigma^2_{\alpha,\beta}$ could likely be done. And further, that there is a slight assumption that the error is normally distributed (just for determining $\sigma^2_{\alpha,\beta}$); though due to the error in the approximation as a whole, the slight assumption should have minimal effect. That is, the actual error need not be normally distributed for the application of the methodology.

### Why error improvement is sensitive to the difference between $Var(\epsilon^\alpha)$ and $Var(O^\alpha - O^\beta)$

For if $Var(O^\alpha - O^\beta)$ is large compared to $Var(\epsilon^\alpha)$, the goal variance $Var(T^\alpha - O^\beta)$ [which equals $Var(O^\alpha - O^\beta) - Var(\epsilon^\alpha)$] will be approximately the same as the variance from the raw localizations $Var(O^\alpha - O^\beta)$, and hence there is no readout to guide the methodology (see the section above).

**Algorithm 1** Method to Maximize Likelihood

See main text for more variable specifics, equations, and definitions<br>
$std(\epsilon^{\alpha})$ is the estimated localization error, std is standard deviation<br>
N is the number of traces<br>
Start with the adjusted positions of locus $\alpha$ equal to observed locations: $A^{\alpha} = O^{\alpha} = (O_1^{\alpha}, O_2^{\alpha}, \ldots O_N^{\alpha})$<br>
BestLik=$CalcLik(A^{\alpha})$ %using equation 3 of main text calculate likelihood for new adjusted positions<br>
epoch=1<br>
**while** $epoch < N \times 5$, **do** %make sure we go until maximized<br>
    $Temp^{\alpha} = A^{\alpha}$ %set temp adjusted values to best performing adjusted locations<br>
    $n = randint(N)$ %randomly pick integer less than N+1 %the number of traces<br>
    $v = randchoose(-std(\epsilon^{\alpha})/20, std(\epsilon^{\alpha})/20)$ %randomly choose one of the two values to adjust by<br>
    $Temp_n^{\alpha} = Temp_n^{\alpha} + v$ %adjust the position of the random localization by small amount<br>
    TempLik=$CalcLik(Temp^{\alpha})$ % calculate likelihood for new adjusted positions<br>
    epoch=epoch+1<br>
    **if** $TempLik > BestLik$ **then**:<br>
        $A^{\alpha} = Temp^{\alpha}$ %Store the new best adjusted positions<br>
        $BestLik = TempLik$ %Store the new best likelihood<br>
        epoch=1<br>
    **end if**<br>
**end while**

**Algorithm 2** Simple algorithm to approximate $\sigma^2_{\alpha,\beta}$

$Array = ()$ #is an empty array<br>
$std(\epsilon^{\alpha})$ is the estimated localization error, std is standard deviation<br>
**for** $epoch$ in $1, \ldots, 500$ **do** #approx using 500 datapoints<br>
    $\epsilon^{sim}$ = array of normally distributed random numbers with a std=$std(\epsilon^{\alpha})$<br>
    $Temp^{\alpha,\beta} = O^{\alpha} - O^{\beta} + \epsilon^{sim}$ #set temp adjusted values<br>
    Append $(Var(Temp^{\alpha,\beta}) - std(\epsilon^{\alpha})^2)$ to $Array$<br>
**end for**<br>
$\sigma^2_{\alpha,\beta}$=$Var(Array)$

### Continued: Verifying resolution improvement with realistic simulation data

We further tested our methodology utilizing the chromatin tracing data of Bintu et al., that quantified the locations of $\approx$ 80 adjacent 30kb segments on a region of chromosome 21 in HCT-116 cells. We selected traces where nearly every locus was localized and assigned the raw experimental locations as the true locations for the analysis and computationally introduced varying degrees of localization error; normal error with $\sigma = 100nm$ or $\sigma = 200nm$. We then applied our methodology and quantified the relative error that resulted. We found that the methodology led to $\approx$ 15% improvement in the localization error for the $\sigma = 100nm$ added error and $\approx$ 30% improvement for the $\sigma = 200nm$ added error (Fig. S2A). Again, clearly showing that the degree of improvement depends on the degree of localization error relative to the distances between the loci; that is, the smaller the distances between loci and the larger the localization error, the better the performance of the methodology.

Lastly, we quantified the performance of the methodology with varying experimental parameters (Fig S2), showing that most existing chromatin tracing datasets meet these standards. We first varied the number of loci included within the analysis (Fig. S2B). Note the loci were always adjacent to each other. Interestingly, we found that with even just three loci the relative error was decreased by 20%, suggesting the use of the methodology with live-cell microscopy data. We believe a central reason why there are diminishing returns when including more loci is because the distances between the loci dramatically increase (in a nonlinear manner) the further away the loci are — that is, if the closer loci were added in last, there would be a dramatic jump in error reduction. Second, we varied the detection efficiency (Fig. S2C). As one would expect the lower the detection efficiency the worse the performance. This result can be thought of as the lower the number of loci, the worse the performance; if a locus is not detected in a particular trace, it will not contribute to improving the localization error in that trace. Third, we investigated the relative error improvement as a function of the degree of localization error guess — the main parameter needed to run the analysis provided by the user, $STD(\epsilon^{\ell})$ (Fig. S2D). For the analyzed dataset with the true $STD(\epsilon^{\ell}) = 200nm$ we varied the different guesses ($\sigma$) from 400nm to 50nm, in every case there was an improvement in the relative error, but if the guess overestimated the true degree of localization error by a huge amount (400nm) some loci did show a worse localization error, still a clear majority showed significant improvements. It is clear that the best improvement was seen when the guess approximated the true value of $STD(\epsilon^{\ell}) = 200nm$, and that no loci showed worse error when underestimating the amount of localization error. Note, we extensively discuss ways to approximate the degree of localization error throughout this Supporting Material. Finally, we varied the number of traces included with the application of the methodology (Fig. S2D), we found that once the number of traces were above 200 the relative error approached a steady value; that is, the number of traces should be at least 300 or higher (at least for this system). Clearly, simple simulations like these may be useful in determining the application of the methodology on a particular experiment.

### Quantifying Localization Error of Initial Localization and Re-Imaged Localization for Chromatin Tracing Data

A straightforward way to quantify the localization error in chromatin tracing data is to re-image the loci and to then quantify the displacement between the two. However, because the re-imaging condition is fundamentally different, the signal between the two is likely different; and therefore, the localization error could also be different. Here we show the methodology we used to determine the 'localization error' of the initially imaged loci and the re-imaged loci. The approach can be easily applied to non-chromatin tracing data. Again, note, here we define the 'localization error' as the standard deviation of the random variable $\epsilon$; the error added to the true locations. See main text for further information on variables.

We start with a pair of loci that were both re-imaged: the observed locations along a single dimension for the two loci ($\alpha$ & $\beta$) are $O^{\alpha}$ & $O^{\beta}$ for the initial image, and $O2^{\alpha}$ & $O2^{\beta}$ are the re-imaged locations. Just as the main text, the observed locations are the following: $O^{\alpha} = T^{\alpha} + \epsilon^{\alpha}$. However, because we have re-imaged loci in this case we write: $O^{\alpha} = T^{\alpha} + \epsilon 1^{\alpha}$ and $O2^{\alpha} = T^{\alpha} + \epsilon 2^{\alpha}$; where $\epsilon 1$ is the error for the initially imaged localizations and $\epsilon 2$ is the error for the re-imaged localizations.

We can then define the following:

$$Contant^{1} = Var(T^{\alpha} + \epsilon 1^{\alpha} - O^{\beta}) = Var(T^{\alpha} - O^{\beta}) + Var(\epsilon 1^{\alpha}) + 2 \times Cov(\epsilon 1^{\alpha}, T^{\alpha} - O^{\beta})$$

$$Contant^{2} = Var(T^{\alpha} + \epsilon 2^{\alpha} - O^{\beta}) = Var(T^{\alpha} - O^{\beta}) + Var(\epsilon 2^{\alpha}) + 2 \times Cov(\epsilon 2^{\alpha}, T^{\alpha} - O^{\beta})$$

$$Contant^{1} - Contant^{2} = Var(\epsilon 1^{\alpha}) - Var(\epsilon 2^{\alpha}) + 2 \times Cov(\epsilon 1^{\alpha}, T^{\alpha} - O^{\beta}) - 2 \times Cov(\epsilon 2^{\alpha}, T^{\alpha} - O^{\beta})$$

Assuming the error is not correlated with anything: $Contant^{1} - Contant^{2} = Var(\epsilon 1^{\alpha}) - Var(\epsilon 2^{\alpha})$. Similarly, we can calculate:

$$Contant^{3} = Var(T^{\alpha} + \epsilon 1^{\alpha} - T^{\alpha} - \epsilon 2^{\alpha}) = Var(\epsilon 1^{\alpha}) + Var(\epsilon 2^{\alpha}).$$

Then,

$$Contant^{3} + Contant^{1} - Contant^{2} = 2 \times Var(\epsilon 1^{\alpha}).$$

And therefore $Var(\epsilon 1^{\alpha})$ can be found, and $Var(\epsilon 2^{\alpha})$, because all three terms on the left are known empirically. This is what we used to quantify the 'localization error' (STD) in the main text.

Here it is important to note that if multiple loci pairs are available (say 100 loci are imaged), the pair of loci with the smallest mean distance should be used. For example, if the goal is to quantify the error of the first locus ($\epsilon 1^{\alpha=1}$), the second locus to use with the above approach will most likely be ($O^{\beta=2}$). The reason for this is due to the error in quantifying $Contant^{1}$ and $Contant^{2}$; the higher the error within the $Var(T^{\alpha} - O^{\beta})$ calculation, the more it will overshadow the $Var(\epsilon 2^{\alpha})$ and $Var(\epsilon 1^{\alpha})$ terms — overall, the error within the variance should scale with the mean, therefore a pair with a low mean distance should be used.

### Approximating Localization Error Using Different Dimensions

Even if loci were not re-imaged to quantify localization error, it is possible to approximate the resolution at least in comparison; given a few assumptions. As above, with a pair of loci ($\alpha$ & $\beta$), the following can be calculated empirically:

$$Var(OX^{\alpha} - OX^{\beta}) = Var(\epsilon X^{\alpha}) + Var(\epsilon X^{\beta}) + Var(TX^{\alpha} - TX^{\beta})$$
$$Var(OY^{\alpha} - OY^{\beta}) = Var(\epsilon Y^{\alpha}) + Var(\epsilon Y^{\beta}) + Var(TY^{\alpha} - TY^{\beta})$$
$$Var(OZ^{\alpha} - OZ^{\beta}) = Var(\epsilon Z^{\alpha}) + Var(\epsilon Z^{\beta}) + Var(TZ^{\alpha} - TZ^{\beta})$$

where $X, Y, Z$ represent the locations of the loci along the different dimensions; that is, $OZ^{\alpha}$ is the observed locations of loci *alpha* along the $Z$ dimension, etc.

Here we must pause for a parentheses and mention the assumptions explicitly. (1) In order for the following to be valid, we assume that the true variances are the same for each dimension — that is, $Var(TX^{\alpha} - TX^{\beta}) = Var(TY^{\alpha} - TY^{\beta}) = Var(TZ^{\alpha} - TZ^{\beta})$; which may not be true in specific cases. And, (2) we assume the error for each locus is more or less the same (since we want to quantify the general amount of localization error); that is, $Var(\epsilon X^{\alpha}) = Var(\epsilon X^{\beta})$, etc.

Nonetheless, we can define the empirically determined constants $C_{zx}$ (or $C_{zy}$) as the following:

$$C_{zx} = Var(OZ^{\alpha} - OZ^{\beta}) - Var(OX^{\alpha} - OX^{\beta}) = 2 \times Var(\epsilon Z) - 2 \times Var(\epsilon X),$$

the localization error along the $Z$ dimension is then:

$$STD(\epsilon Z) = \sqrt{C_{zx}/2 + STD(\epsilon X)^2},$$

where $STD()$ is the standard deviation. This means, by knowing something about the localization error along one dimension, you can determine the localization error along the other dimension.

#### Quantifying the Worse Case Scenario for Localization Error

For the benefit of the reader, we are going to discuss a brief calculation that we did not heavily utilize within this work but could prove useful when analyzing multi-loci microscopy data. The calculation follows the question:can we get an idea of a higher bound and lower bound on localization errors? To gain insight toward half of this question, we start by considering the worse case scenario, only using the distances between adjacent loci, and then comparing similar systems to approximate the error to approximate the lower bound (see below).

We quantified the displacements between adjacent loci for each dimension and quantified the variance; that is: $Var(O^k - O^{k+1})$. Expanding with the error (See above and main text):

$$Var(O^k - O^{k+1}) = Var(T^k - T^{k+1}) + Var(\epsilon^k) + Var(\epsilon^{k+1}).$$

We then assume that the error for both loci are the same ($STD(\epsilon^k) = STD(\epsilon^{k+1})$), and set $Var(T^k - T^{k+1}) = 0$; note, that the previous, in reality, will not be zero. This is what we mean by worse case scenario — for the quantified localization error has to be less given that the term is not zero.

The worst case scenario for the localization error is:

$$STD(\epsilon) \approx \sqrt{Var(O^k - O^{k+1})/2} \approx \sqrt{Var(\epsilon^k)/2 + Var(\epsilon^{k+1})/2} \approx \sqrt{Var(\epsilon)}$$

This serves as a good sanity check to make sure the localization errors are not overestimated. It should be noted, that the better the genomic resolution, the better the worse case scenario will approximate the true localization error.

### Specifics for using the methodology on the Huang et al. dataset

To apply the methodology to the dataset of Huang et al. (4) we needed to estimate the localization error of their experiments. Problematically, they did not re-image any of the loci, preventing us for quantifying the localization error directly. We therefore had to take several different approaches to approximate these values.

First, because the genomic resolution was so fine (5kb genomic resolution), we first sought to determine the worst case scenario for the localization errors. The 'worst case scenario' approach is described in depth above within the Supporting Material. This resulted in a localization error centered around 100nm for the $x, y$ dimensions and 125nm for the $z$ dimension. The probability distributions of the 'worse case scenario' calculations are shown in Fig. S3A. Again, because the genomic resolution was so fine, this calculation likely approximates the true localization error, but the actual value is likely slightly lower.

To explore further, we sought to compare the Huang et al. dataset to a dataset where the localization error had been quantified. To do this we calculated the distances between neighboring loci for the Su et al., dataset (9) and the Huang et al. dataset, and compared the distributions (Fig. S3B). Intuitively, the proportion of the smallest distances are limited by the localization error, and therefore we should be able to compare the two to some extent. However, the comparison is made complicated due to the greatly different genomic resolutions, 50kb for the Su et al. dataset. Still, naturally we found the 5kb genomic resolution of Huang et al. had the highest proportion of smaller distances — we believe this to be primarily due to the polymer nature of chromatin. To make the comparison more 'fair' but still biased toward low localization errors for Huang et al., we quantified the distances between 'neighboring loci' within Huang et al. but this time artificially increased the genomic resolution to 35kb; as expected, the proportion of smaller distances decreased. Here we note that this is an approximation in terms of the comparison, as the Su et al. dataset positions are the centroid positions of 50kb segments, while the pseudo 35kb dataset uses the centroids of 5kb segments located 35kb away from each other. When compared to the raw Su et al. distances, the proportion of low distances was higher for the 50kb genomic resolution dataset. Note, that these distances were only quantified along a single dimension for all datasets ($X$). In order to 'match' the pseudo 35kb data, we had to computationally introduce more localization error; incorporating normally distributed error with a standard deviation of 50nm. Only then were the distances of the 50kb dataset comparable. Because the Su et al. dataset already had a localization error of std=50nm along the $X$ dimension, the computationally incorporated error raises the std of the localization error to approximately 70nm.

Taken together, we decided to go with a likely underestimation of the localization error along the $X, Y$ dimension, using $STD(\epsilon) = 70nm$. This way we do not accidentally incorporate artifacts by removing too much variability. The reasons why this is likely an underestimation is because we used a pseudo genomic resolution of 35kb instead of 50kb, due to the polymer nature of the data this comparison should result in an underestimation of the error. Further, the fact that the genomic resolution is so fine with the original Huang et al. dataset, points toward the 'worse case scenario' approaching the true localization error; along the $X$ dimension 70nm is clearly on the low end of the spectrum. With the estimation for the $X, Y$ dimensions we were then able to estimate the localization error along the $Z$ dimension at approximately 100nm; see the section 'approximating localization error using different dimensions.'

### Additional Filtering of Chromatin Tracing Data to Eliminate Extreme Off Target Error

For all of the chromatin tracing data probed, we performed an additional filtering step to eliminate localizations that overwhelmingly result from off target error. Note, without this step we found that the amount of localization error (in terms of standard deviations quantified from re-imaged loci) was amazingly large; often above 400 nm (data not shown) — one of the observations that directed us to investigate the error further. Here we should further state that if one is concerned that the following process may remove actual localizations, artificially reducing the variability of the data for the sake of the application of the methodology, one could simply apply the methodology and then 'add back' the filtered out localizations. Nonetheless, to explain the logic of the filtering procedure we first performed a simple analysis and discuss the results; we show an illustration for guidance (Fig. S4). In the following sections, we also discuss the specifics of how we applied this filtering to each dataset and how each dataset supports the approach, and further why the error improvement with the application of the methodology is underestimated with actual experimental data.

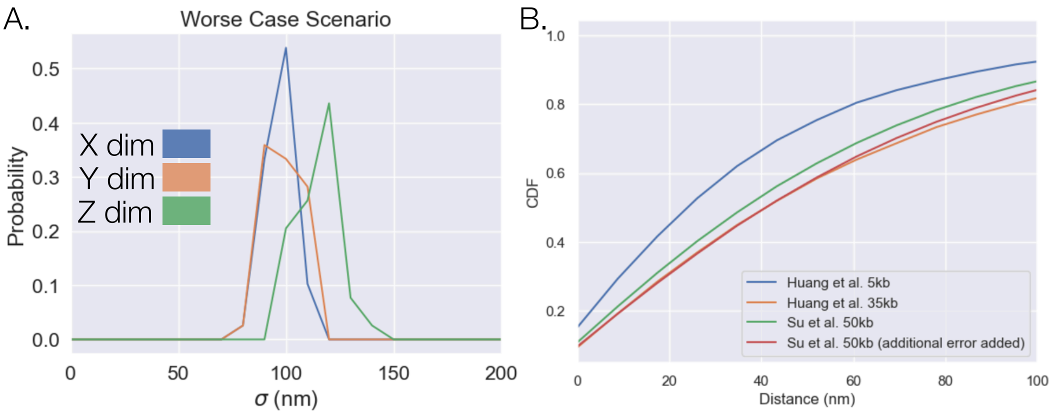


Figure S3: *Approximating the localization error of Huang et al.:* A) The 'worst case scenario' quantified for the different dimensions of the raw dataset. B) The cumulative distribution for the distances between neighboring loci at different genomic resolutions. These distances were only quantified along the $X$ dimension.

The approach relies upon a metric we call the *Minimum Distance to Neighboring Loci* (Fig. S4); for each localization (of locus $i$, Fig. S4), we calculated the distances to each of the neighboring loci (of loci $i+1$ and $i-1$) and kept whatever distance was lower for each individual trace. We did this for both the initially imaged loci and the re-imaged loci. The idea is to use this metric to filter out localizations with extreme amounts of error, which appear to be the result of off targeting. Put another way, filter out localizations that have a large *Minimum Distance to Neighboring Loci* for, as we demonstrate, these were the localizations with extreme amounts of error due to off targeting.

To understand the relationship to the localization error of individual localizations, we plotted that which is illustrated in Fig. S4; this was done using the re-imaged loci of real experimental data. More specifically, the distance between the initially imaged localization and the re-imaged localization is shown on the plot with color (nm); red illustrates a large displacement error and blue a low displacement error. The *min distance to neighboring loci* for the initial localization is shown on the x axis (nm), while that for the re-imaged loci is shown on the y-axis (nm). Note, the data in the colored plot is the 2kb resolution chromatin tracing data from Mateo et al. 2019 using only the Z-dimension (7). Similar plots were seen for all datasets used within the main text and are described in the following sections.

The plots have three regions of interest; these are emphasized in Fig. S4. Region 1 shows that once the *Minimum Distance to Neighboring Loci* gets above a certain point, about 400nm for this particular example, the displacement error goes extremely high; the same was seen along the y-axis for region 2. The fact that the almost all of the localizations with extreme amounts of displacement error fall into these two regions suggests that the datapoints result from either the initial localization or the re-imaged localization targeting something other than the intended loci (off target) — this would explain the huge amounts of "error," as they are not targeting the same locus. Therefore, to eliminate localizations not targeting the proper locus one could eliminate localizations that have a large *Minimum Distance to Neighboring Loci*.

However, what about region 3? One could argue that many localizations with large *Minimum Distance to Neighboring Loci* were actually targeting the proper target because the displacement error is low, aka the localizations report on the correct target. Put another way, the locations within region 3 show small displacement error even with large *Minimum Distance to Neighboring Loci*, suggesting that they are reporting on the same target. We ultimately found that the localizations within region 3 were also due to off target localizations; we discuss the localizations within region 3 extensively throughout the following subsections, as demonstrating this requires comparisons between different chromatin tracing datasets. Overall, when we applied this to each experimental dataset the new quantified localization errors (in terms of standard deviations) approached reasonable values ranging from 40nm to 150nm.

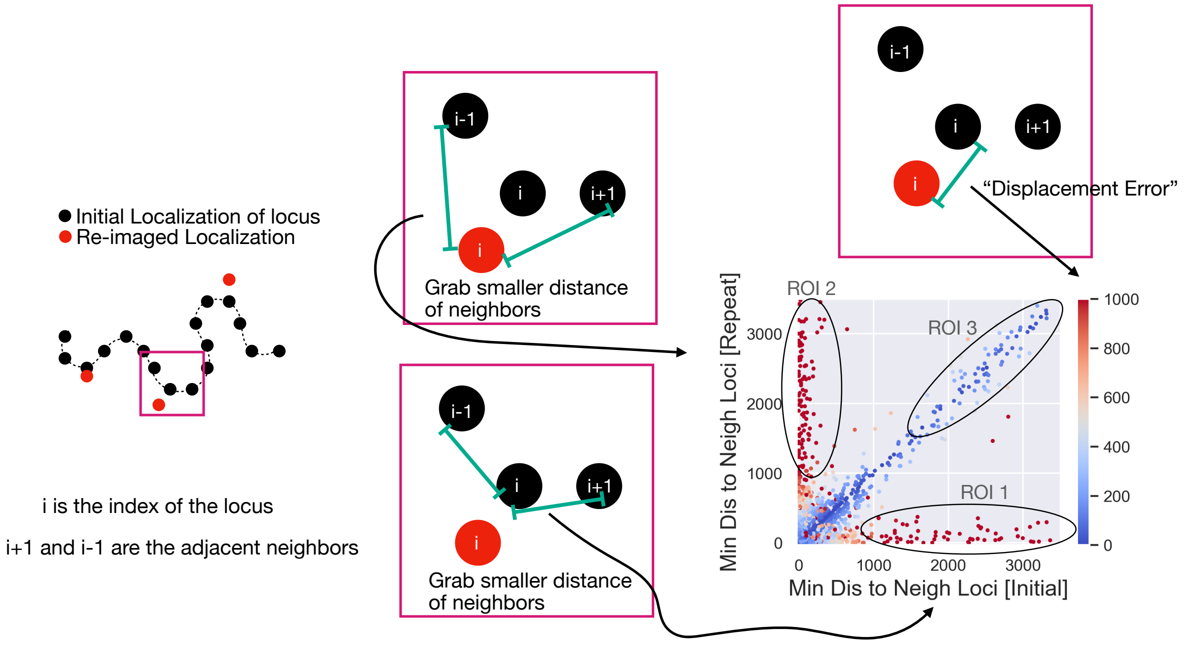


Figure S4: *Illustration of 'displacement error' and 'minimum distance to neighboring loci':* On the far left is an example trace with the initially imaged loci in black and those that were re-imaged in red. The index of the loci are explained within the figure. The center two boxes illustrate the definition of 'Min dis to neigh loci' using either the initial imaged loci (bottom box) or the re-imaged loci (top box). The boxes illustrate that the distances to the neighboring loci are calculated and then the smaller one is taken; using either the initially imaged or the re-imaged. Each dot on the plot on the bottom right is from a single trace. The color of each dot shows the displacement error, which is calculated as the distance between the initial and re-imaged localizations for each trace. The 'displacement error' is essentially a proxy for the localization error of individual localizations, and the calculation is illustrated with the top right box. In the plot we also highlight three regions of interest (ROI), and each of these are described within the text of the Supporting Material.

#### Why Error Improvement from Methodology was Underestimated Empirically

Continuing from the section above, we found that the localizations with a high *Minimum Distance to Neighboring Loci* were primarily the result of error; we believe primarily due to mis-targeting (see below for further discussion). Regions 1 and 2 clearly show that the repeat localization and the initial localization do not report on the same target when the *Minimum Distance to Neighboring Loci* is high.

But why do we claim that the error improvement from the methodology is underestimated with the actual experimental data? The approach we used to filter out these off target localizations was setting a threshold and removing all localizations with a *Minimum Distance to Neighboring Loci* higher than the threshold. Still, some off target localizations undoubtably remain. Considering that the localization error improvement is quantified by measuring the distances between the initial and re-imaged loci; if they are reporting on different targets the distance between the two will not get smaller if the localization error is minimized. Hence, the off target localizations within experimental chromatin tracing data will cause the error improvement from the methodology to be underestimated.

#### Filtering Justification of Chromatin Tracing Data of Su et al. (Addressing ROI 3)

We found that the dataset with the most re-imaged loci was the sequential imaging data of Su et al., and we show a careful error interrogation of this dataset in Fig. S5. Also, given that there were other chromatin tracing datasets within IMR90 cells (1), a comparison of the datasets suggested that the localizations within the previously mentioned region 3 should be filtered out.

To get an overall idea of the localization error within this chromatin tracing dataset, we quantified the 'displacement error' for each dimension; that is, we quantified the distance between the initially imaged localizations and the re-imaged localizations. The results of this analysis are shown in Fig. S5A and B., on a linear scale and a log scale respectively. The $X$ and $Y$ dim are shown in orange and green while the $Z$ dimension is shown in blue. We also show the displacement error which results from normally distributed error with a standard deviation of 100nm in black. We found that upon first glance, on a linear scale the displacement error along the $X$ and $Y$ dimension was lower than that of the $Z$ dimension (as one would expect), and the displacement error along the $Z$ dimension was similar to the simulation. Interestingly though, when shown with a log scale on the y-axis, the displacement error showed an extremely long tail; with the $X$ and $Y$ dimensions performing even worse than the $Z$ dimension.

We next sought to understand whether we could filter out the localizations using the *Minimum Distance to Neighboring Loci*. Therefore, we sought to determine whether the localizations with the extreme amount of error were also the localizations with the large *Minimum Distance to Neighboring Loci*. We generated the plots described in the section above, showing the relationships between the *Minimum Distance to Neighboring Loci* and the displacement errors (Fig. S4). Along all three dimensions, we observed all three previously discussed regions of interest. Regions 1 and 2, clearly suggest using a *Minimum Distance to Neighboring Loci* cutoff of around 500nm; the displacement error increased drastically once the *Minimum Distance to Neighboring Loci* got above this value, at least for regions 1 and 2 for all dimensions. Still, we found that the localizations in region 3 had a low displacement error, suggesting that some of the localizations with large *Minimum Distance to Neighboring Loci* were 'good.' Put another way, again, there was the possibility that these localizations were the result of the large degree of heterogeneity in chromatin structure and should therefore be kept.

To probe further, we compared the *Minimum Distance to Neighboring Loci* in an orthogonal chromatin tracing dataset to the Su et al. dataset, ultimately showing that the vast majority of the localizations in region 3 should be discarded. The work of Bintu et al. quantified multiple chromatin regions within IMR90 cells (the same cell type as that of Su et al.) at a genomic resolution of 30kb (Fig. S6A and C). Because the genomic resolutions of the datasets were not the same, we skipped over loci within the Bintu et al. dataset to make the genomic resolution 90kb; the larger genomic resolution and the polymer nature of chromatin would make the *Minimum Distance to Neighboring Loci* even larger. We show the cumulative distributions of all *Minimum Distance to Neighboring Loci* in Fig. S6B and D. When we compared these CDFs with those from the sequential dataset of Su et al. (Fig. S6F and Fig. S5D), we found them comparable up to approximately 500nm; we observed 98% and 96% of the *Minimum Distance to Neighboring Loci* were below 500nm for the dataset of Bintu et al., while about 96% were below 500nm for Su et al. However, we found that as the *Minimum Distance to Neighboring Loci* increased above 500nm the more the Bintu et al. dataset deviated; there were absolutely no values above 1000nm for Bintu et al., but about 1% for Su et al. Showing a disagreement between the two datasets, suggesting that the vast majority of localizations with high *Minimum Distance to Neighboring Loci* are incorrect and should be removed (Further evidence provided with the Mateo et al. dataset below).

We therefore eliminated localizations from the dataset of Su et al. with a *Minimum Distance to Neighboring Loci* value greater than 500nm. For the dataset of Su et al. we initially selected for traces where the percentage of detected loci was above 90% (Fig. S5E, dashed line). After, eliminating localizations with high *Minimum Distance to Neighboring Loci*, this decreased the % of loci detected by about 10% (Fig. S5E, solid line).

Note, further evidence that the localizations in region 3 should be discarded are further supported in the following section with the error analysis of the 2kb chromatin tracing data of Mateo et al.

#### Filtering Justification for the Chromatin Tracing Data of Mateo et al. (Addressing ROI 3 Further)

Within the chromatin tracing data of Mateo et al. there are multiple loci re-imaged for both the 2kb chromatin tracing dataset and the 10kb chromatin tracing datasets. In this section, we first discuss the localization error and the justification for filtering localizations for the 2kb dataset — as this dataset more clearly supports the argument that the localizations in region 3 result from off target localizations. We then go on to discuss the error and the filtering of the 10kb dataset. Note, the quantification and interpretation for this dataset is similar to that of Su et al. dataset described above.

Again we quantified the displacement error individually for each dimension of the 2kb resolution tracing data. We found that for this dataset the number of re-imaged loci was 6, lower than that within Su et al. The displacement

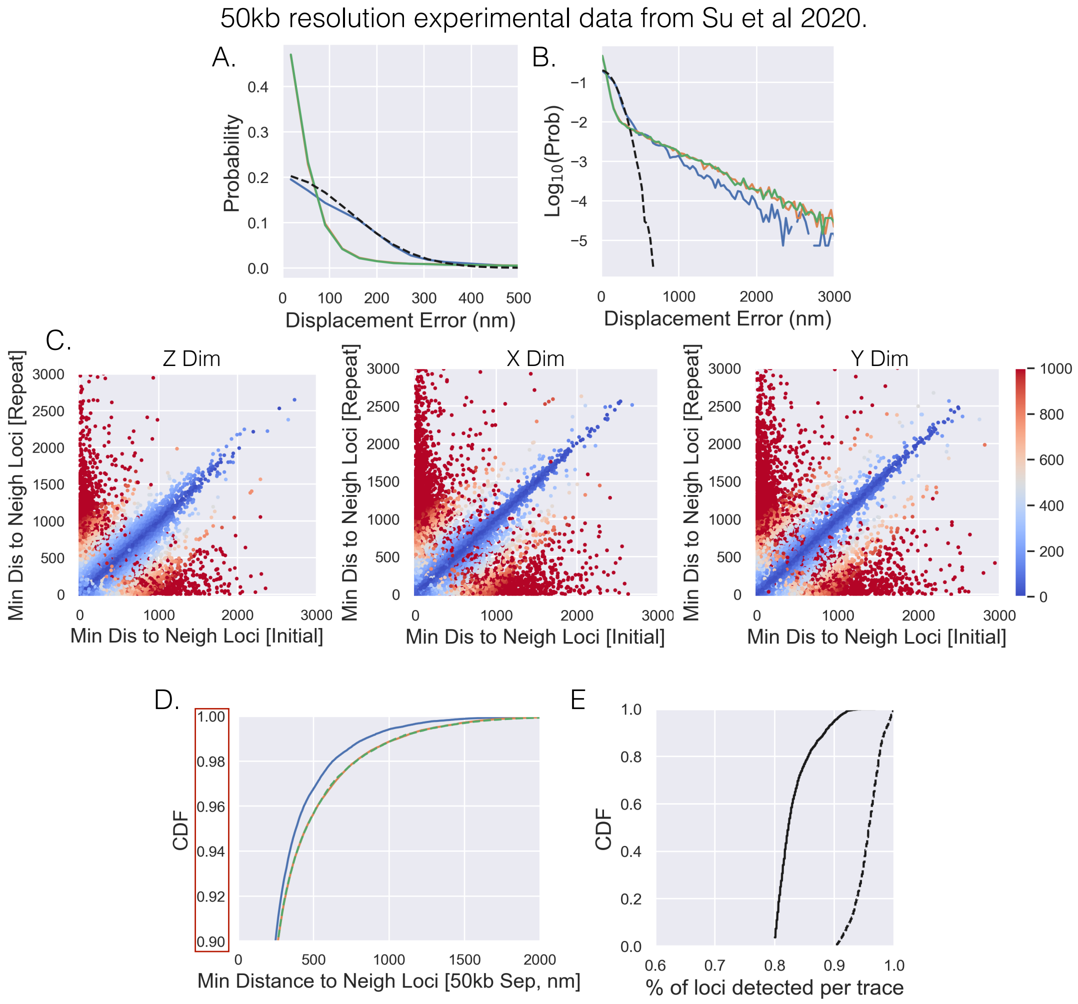


Figure S5: *Investigating Su et al chromatin tracing data and localization error:* A) The probability distributions for the displacement error (the distance between initially imaged loci and re-imaged loci) for each of the imaging dimensions; blue is $Z$ dimensions and green and yellow are $X$ and $Y$. We also show with simulation the probability distribution for loci with a standard deviation equal to 100nm; black dashed line. B) The same as 'A' but with a log scale to highlight the long tail of the distributions. C) The minimum distance to a neighboring loci for the re-imaged localization vs. the minimum distance to a neighboring loci for the initially imaged localization, where the color shows the displacement error along the labelled dimension. Please note the metrics are illustrated in Fig. S4. D) The cumulative distribution for all minimum distances to neighboring loci for each of the individual dimensions, highlighting that a small proportion of localizations have high minimum distances to neighboring loci. E) The cumulative distribution for the detection efficiency of individual traces. The dashed line is for the traces before eliminating localizations with high minimum distances to neighboring loci while the solid line is after.

error results are shown in Fig. S7A and B., on a linear scale and a log scale respectively. [Again, the $X$ and $Y$ dim are shown in orange and green while the $Z$ dimension is shown in blue.] We again show the displacement error that results from normally distributed localization error with a standard deviation of 100nm, shown as a black dashed line. Similar to the dataset of Su et al., the localization errors looked as expected when shown on a linear scale (Fig. S7A), and showed displacements consistent with a localization error better or equivalent to that of the simulation. But again, the displacement error showed a long tail extending well out to 1000nm when on a log scale (Fig. S7B), suggesting an extreme amount of error for some localizations.

To investigate if we could filter out the localizations with extreme amounts of error given *Minimum Distance to*

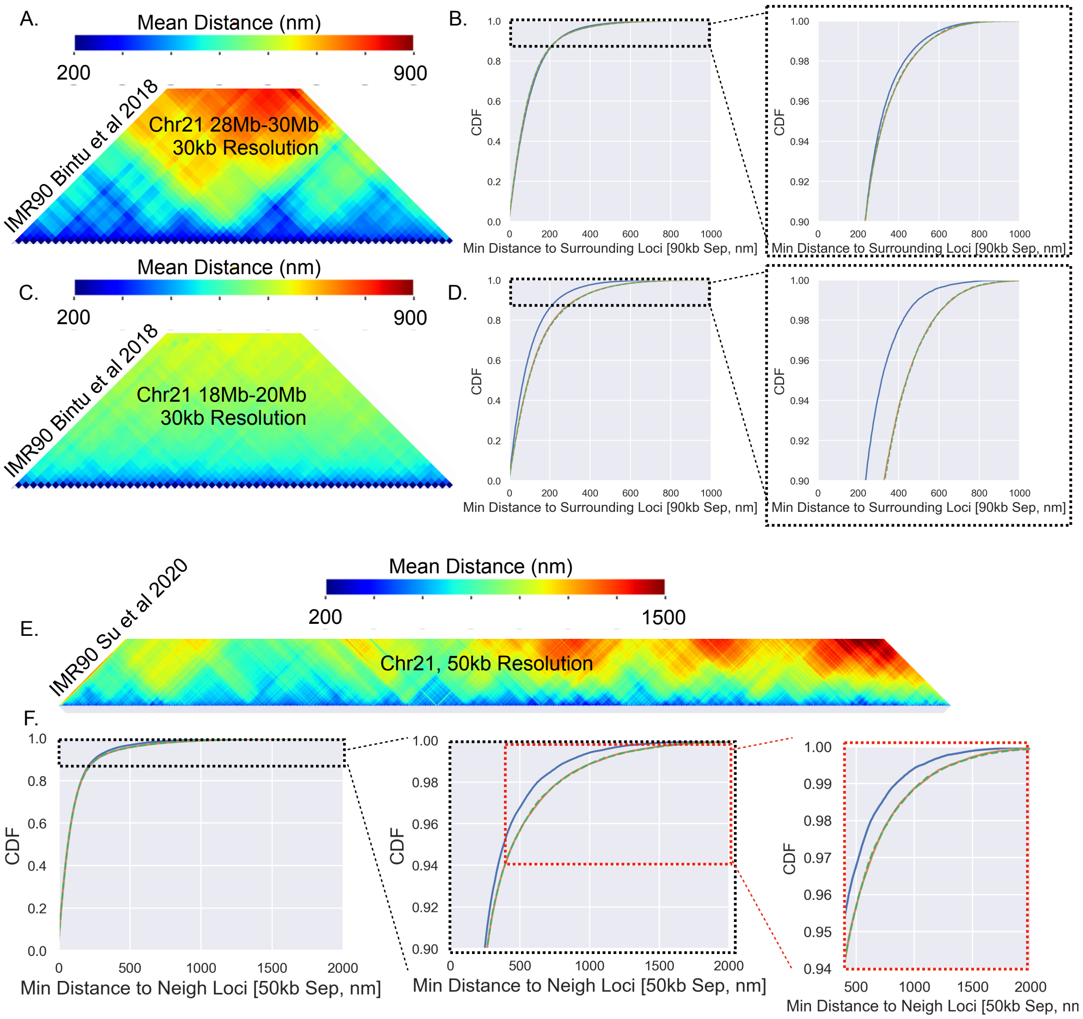


Figure S6: *Comparing multiple chromatin tracing datasets suggests that localizations with high minimum distances to neighboring loci are off target:* A+C+E) The mean distances between loci for the specified chromatin tracing dataset. B+D+F) The cumulative distribution function for the different chromatin tracing datasets with zoom ins highlighting the largest minimum distances to neighboring loci. Note that for 'B' and 'D' we artificially made the genomic resolution 90kb so that they would be more comparable with 'F.' That is, the larger the genomic resolution the larger the distances between neighboring loci.

*Neighboring Loci* threshold, we sought to determine whether the localizations with the extreme amount of error were also the localizations with the large *Minimum Distance to Neighboring Loci*. The *Minimum Distance to Neighboring Loci* for the initial localization and the repeat localization with their corresponding displacement error are shown in Fig. S7C. For all three dimensions we saw the three regions of interest as described in the above sections. We found that a *Minimum Distance to Neighboring Loci* threshold of around 400nm for all three dimensions would eliminate the vast majority of localizations with extreme displacement error for all dimensions in regions 1 and 2.

What about region 3 with this dataset? That is, were the localizations with high *Minimum Distance to Neighboring Loci* and low displacement error targeting the correct loci? We noticed a great difference between the different dimensions, with the *Minimum Distance to Neighboring Loci* along the $Z$ dimension in region 3 extending well above 1000nm, going up to around 3000nm, while those along $X$ and $Y$ topped out around 1000nm. If the chromatin organization is symmetric in all three dimensions, the localizations in region 3 of $Z$ must be 'wrong.' Note, here we should state that the fine genomic resolution of this dataset likely makes the application of chromatin tracing more difficult, a likely reason for why that from the $Z$ dimension is so different from the $X$ and the $Y$ dimension. This result again provides strong support that the localizations with high *Minimum Distance to Neighboring Loci*, even in region 3, should be removed.

We therefore removed localizations with a *Minimum Distance to Neighboring Loci* greater than 400nm. Note, we reasoned that the slightly lower *Minimum Distance to Neighboring Loci* threshold (when compared to that applied to the Su et al. dataset) was justified given the genomic resolution here was 2kb compared to 50kb — naturally, the *Minimum Distance to Neighboring Loci* should be higher for larger genomic distances. When quantifying this dataset, we only utilized traces where the % of loci detected was above 75%, the traces had a median detection around 85% (Fig. S7E, dashed line). When we filtered out the localizations with a *Minimum Distance to Neighboring Loci* above 400nm, the median detection fell to about 60% (Fig. S7E, solid line).

Yet, given that the chromatin tracing was done in fly embryos, the assumption that the chromatin structure was symmetric in all dimensions may not hold. We addressed this with a further comparison with the 10kb resolution data, which also suggests that the localizations with large *Minimum Distance to Neighboring Loci* are the result of error. Just as before, we quantified the displacement error for each dimension and show the results in Fig. S8A and B. Reproducibly, we saw that the displacement error distributions had long tails when shown on a log scale. We then generated the previously described plots and found all three regions of interest (Fig. S8C), again supporting the idea of using a *Minimum Distance to Neighboring Loci* threshold to remove problematic localizations. In regards to the comparison, due to the polymer nature of chromatin the 10kb resolution data should have larger *Minimum Distance to Neighboring Loci* when compared to the 2kb resolution data; generally, the larger the genomic distance the larger the physical distance between neighboring loci. Appealingly for the filtering approach, we found that the quantification for the $Z$ dim of the 10kb data had *Minimum Distance to Neighboring Loci* values top out around 1000nm (Fig. S8C); hence, the vast majority of the localizations within the 2kb dataset with high *Minimum Distance to Neighboring Loci* values are the result of error (even for region 3) — as supported by all of the analyses above. Another line of reasoning is that the 10kb data demonstrates that the chromatin structure does appear to be more or less symmetric (at least when comparing these metrics); consequently, the arguments made with the 2kb stand regarding region 3.

#### Specifics for Applying the Methodology to the Bintu et al. dataset

To apply the methodology to the dataset of Bintu et al. we needed an estimation of the localization error. Within the work of Bintu et al. they utilized two microscopy techniques when collecting the chromatin tracing data, STORM and simple epiflourence microscopy. Here it should be noted when using the later microscopy technique, individual diffraction limited spots were still fit enabling superresolution microscopy; still, the STORM image data undoubtedly has a superior resolution. Problematically, loci were only re-imaged for the STORM experiments, finding a localization error of 50nm for each dimension. Even though the epi-images have a worse localization error, we decided to use an underestimation of the localization error, with an estimation of 50nm for each dimension. Even though the error improvement from the methodology will be limited in this case, this way every loci will likely show an improvement in localization error with the methodology. For if too high an estimate of the localization error is used, a low proportion of the loci could actually show a worse localization error after the application of the methodology (Fig. S2). Consequently, we decided to 'play it safe' and underestimate the localization error for this analysis.

### Calculating the expected independent higher order contacts

The calculation we used to quantify the expected value for higher order contact frequencies is illustrated with the following example. To quantify the expected 3-way contact frequency between the loci $\alpha, \beta$, and $\gamma$, we first quantify the 2-way contact-frequencies between each pair $(P(\alpha : \beta), P(\alpha : \gamma), P(\beta : \gamma))$; again, this is the proportion of time, or the probability that two loci are within a certain distance of each other. For independent events, we simply multiply the probabilities, this is what we refer to within the main text as the expected independent 3-way contact frequency.

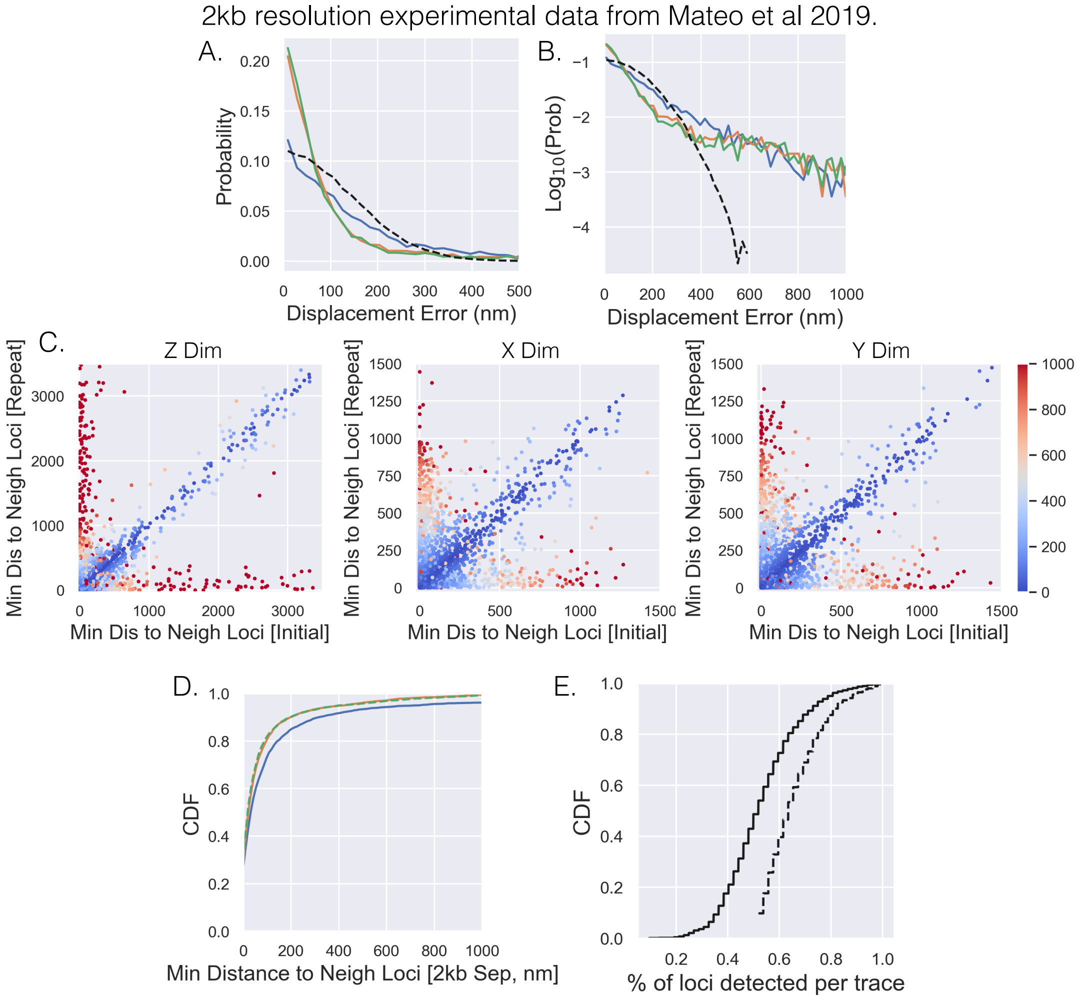


Figure S7: *Investigating 2kb chromatin tracing data and localization error of Mateo et al:* The same analysis as in Fig. S5 but with the 2kb chromatin tracing data of Mateo et al.

### Example for cooperativity as a buffer for multi-way contacts

We illustrate how the cooperativity could act as a buffer with an example. Hypothetically, say with loop extrusion and no cooperatively an independent 3-way contact frequency of .2 is seen, but with cooperatively the 3-way contact frequency of $\approx$ .3 is observed (Fig. 5D, see main figure for guidance). Now, say with the elimination of loop extrusion, the 2-way contact frequencies are dramatically decreased leading to an independent 3-way contact frequency of .05 for the same three loci, again with cooperatively the 3-way contact frequency ends up being .15 (Fig. 5D). In terms of relative changes, the multi-way contacts with cooperatively lead to a 50% decrease, while without cooperativity there was a 75% decrease — which could be significant depending upon the exact relation between contact frequencies and transcription (2, 3, 5, 6). Taken with studies that have shown 'relatively' small changes in transcription activity with the elimination of loop extrusion, perhaps this 'buffering' mechanism is contributing (8).

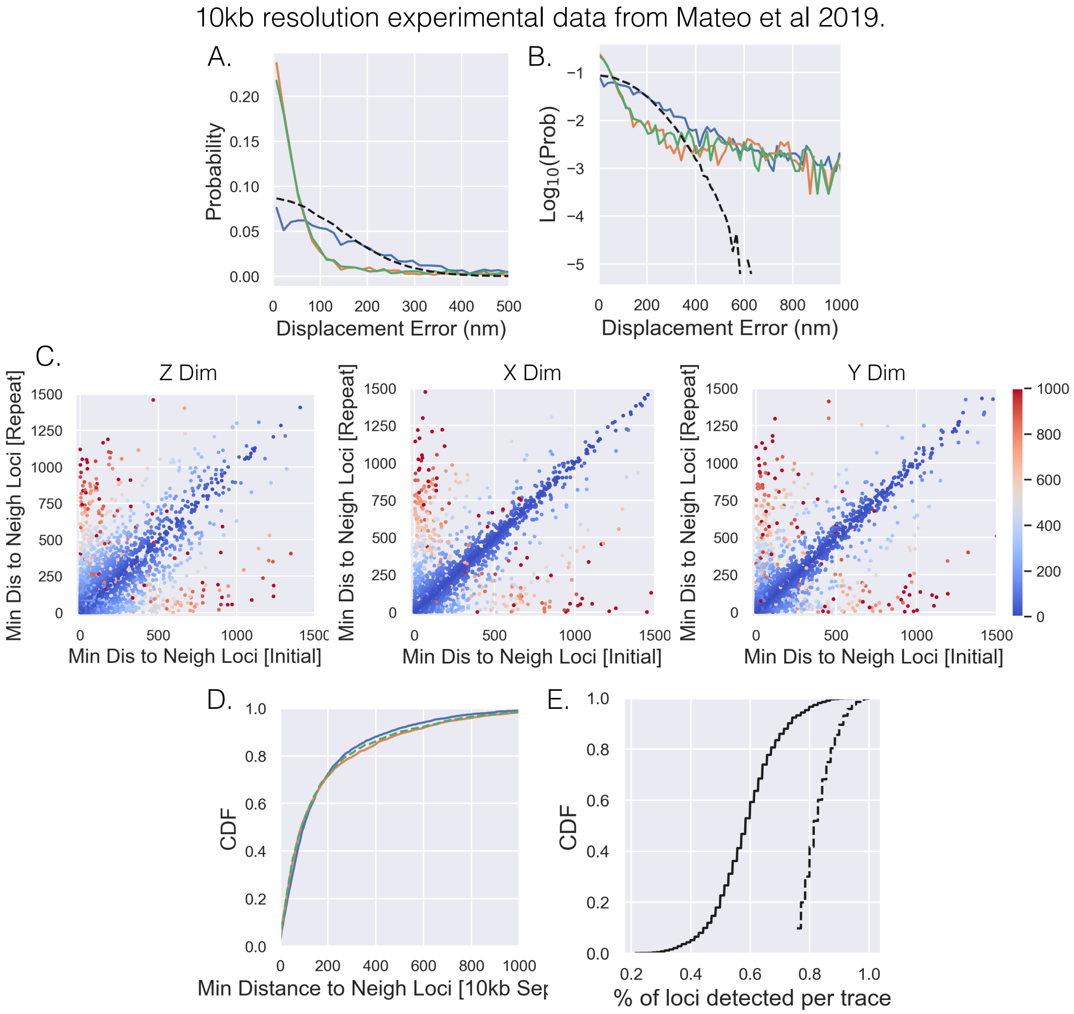


Figure S8: *Investigating 10kb chromatin tracing data and localization error of Mateo et al:* The same analysis as in Fig. S5 but with the 10kb chromatin tracing data of Mateo et al. Note, here the number of loci that were re-imaged was two, which is why the amount of data is much less.

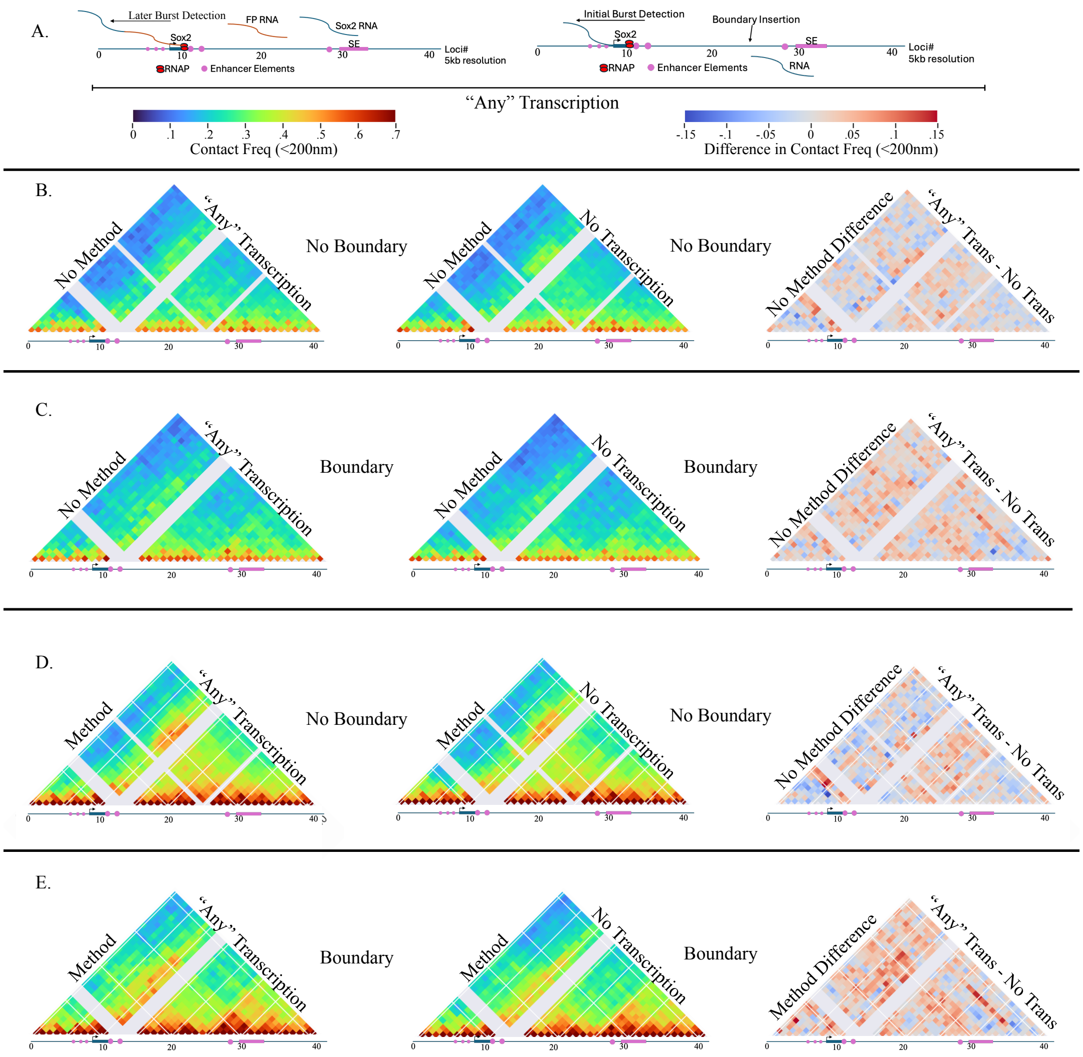


Figure S9: *The linkage between Sox2 transcription and the organization of chromatin:* A) An illustration of the *Sox2* region with the two types of FISH signal that Bing et al. used. In this experimental system, a fluorescent protein (FP) sequence was placed after the *Sox2* gene and different probes targeted the *Sox2* RNA or the FP RNA. If both RNAs were detected at a single allele, we refer to that as a 'later burst detection.' If only the *Sox2* RNA was detected at an allele, we call that situation an initial burst, for the RNAP would not have traveled as far after initiation. The 'any transcription' category is for an allele where either a later burst or an initial burst was detected. B) The contact frequency maps for the no boundary condition using the raw data, comparing traces with any kind of transcription (left) to the no transcription frequencies (middle). The difference between the two are shown on the far right. C) The same as *B* but using the raw data boundary condition data. D) The same as *B* but using the methodology. E) The same as *C* but using the methodology.
